# Nigrostriatal Dopamine Signals Sequence-Specific Action-Outcome Prediction Errors

**DOI:** 10.1101/2021.01.25.428032

**Authors:** Nick G. Hollon, Elora W. Williams, Christopher D. Howard, Hao Li, Tavish I. Traut, Xin Jin

## Abstract

Dopamine has been suggested to encode cue-reward prediction errors during Pavlovian conditioning. While this theory has been widely applied to reinforcement learning concerning instrumental actions, whether dopamine represents action-outcome prediction errors and how it controls sequential behavior remain largely unknown. Here, by training mice to perform optogenetic intracranial self-stimulation, we examined how self-initiated goal-directed behavior influences nigrostriatal dopamine transmission during single as well as sequential instrumental actions. We found that dopamine release evoked by direct optogenetic stimulation was dramatically reduced when delivered as the consequence of the animal’s own action, relative to non-contingent passive stimulation. This action-induced dopamine suppression was specific to the reinforced action, temporally restricted to counteract the expected outcome, and exhibited sequence-selectivity consistent with hierarchical control of sequential behavior. Together these findings demonstrate that nigrostriatal dopamine signals sequence-specific prediction errors in action-outcome associations, with fundamental implications for reinforcement learning and instrumental behavior in health and disease.

## INTRODUCTION

Our brains constantly generate predictions about the world around us (Rao and Ballard, 1999; Keller and Mrsic-Flogel, 2018), particularly regarding the expected consequences of environmental cues or our own actions (Wolpert et al., 1995; Crapse and Sommer, 2008; Schneider et al., 2018; Wurtz, 2018). Indeed, the effects of such expectations have long been recognized when examining the phasic activity of midbrain dopamine neurons following reward-predictive stimuli during Pavlovian conditioning (Fiorillo et al., 2003). Many dopamine neurons signal errors in cued reward prediction (Houk et al., 1995; Montague et al., 1996; Schultz et al., 1997; Cohen et al., 2012; Eshel et al., 2015, 2016; Engelhard et al., 2019), i.e., any change in expectation of future reward or difference between actual versus expected reward predicted by the cues (Sutton and Barto, 2018). These dopaminergic prediction errors are thought to convey a teaching signal that is critical for multiple forms of associative learning across the corticostriatal topography (Yin et al., 2008; Balleine, 2019), spanning both classical Pavlovian stimulus-outcome conditioning (Flagel, et al., 2011; Steinberg et al., 2013; Chang et al., 2016; Saunders et al., 2018; Maes et al., 2020) and the formation of stimulus-response habits (Knowlton et al., 1996; Matsumoto et al., 1999; Faure et al., 2005; Belin and Everitt, 2008; Wang et al., 2011; Kim et al., 2015).

However, the vast majority of previous studies examining dopamine responses primarily have used discrete reward-predictive stimuli (Schultz et al., 1997; Fiorillo et al., 2003; Morris, et al., 2004; Roesch et al., 2007; Flagel, et al., 2011; Hollon et al., 2014; Cohen et al., 2012; Eshel et al., 2015, 2016; Matsumoto and Hikosaka, 2009; Parker et al., 2016; Coddington and Dudman, 2018; Engelhard et al., 2019), whether Pavlovian conditioned stimuli, for which no action is required to earn reward, or explicit discriminative stimuli that essentially instruct an animal how and when to respond to earn reward. Although such explicit cues can exert powerful influences over behavior, far less is known regarding dopamine’s potential roles in and interactions with goal-directed behavior that is self-initiated, self-paced, and guided by instrumental action-outcome associations.

Parallel work also has sparked a renewed focus on dopamine in movement control (Klaus et al., 2019; Coddington and Dudman, 2019). Across multiple recording modalities, several recent studies have reported prominent changes in dopamine activity at the initiation of spontaneous movements ranging from operant action sequences (Jin and Costa 2010; Wassum et al., 2012; Collins et al., 2016; da Silva et al. 2018) to locomotion and brief postural adjustments (Barter et al., 2015; Dodson et al., 2016; Howe and Dombeck, 2016; da Silva et al. 2018; Coddington and Dudman, 2018). However, the extent to which such movements were directed toward any particular goal in the latter studies is unclear, and how self-initiated goal-directed actions influence nigrostriatal dopamine release is largely unknown. Therefore, despite longstanding implication in voluntary movement, motivation, and reinforcement learning, the precise role of dopamine in instrumental action remains poorly understood. Here, we examined how goal-directed action and learned action sequences influence the nigrostriatal dopamine response to the consequence of these actions, in behavioral contexts with minimal overt changes in the animal’s external environment.

## RESULTS

### Suppression of Optogenetically Stimulated Nigrostriatal Dopamine by Goal-Directed Action

Mice expressing channelrhodopsin-2 selectively in their dopamine neurons (See Methods; Zhuang et al., 2005; Madisen et al., 2012) were implanted with a fiber optic over the substantia nigra pars compacta (SNc) for optogenetic stimulation (Sparta et al., 2011) and a carbon-fiber microelectrode (Clark, Sandberg et al., 2010) in the ipsilateral dorsal striatum to record nigrostriatal dopamine transmission using fast-scan cyclic voltammetry (FSCV; Figure 1A, Figure 1—figure supplement 1A-F). The mice were trained in a free-operant optogenetic intracranial self-stimulation (opto-ICSS) task (Figure 1B), in which they learned to press a continuously reinforced “Active” lever to optogenetically stimulate their own dopamine neurons (50 Hz for 1 s) and rarely pressed the non-reinforced “Inactive” lever yielding no outcome (Figure 1C). Therefore, consistent with other recent reports (Rossi et al., 2013; Ilango et al., 2014; Keiflin et al., 2018; Saunders et al., 2018), selective stimulation of SNc dopamine neurons is sufficient to reinforce novel actions.

**Figure 1.**
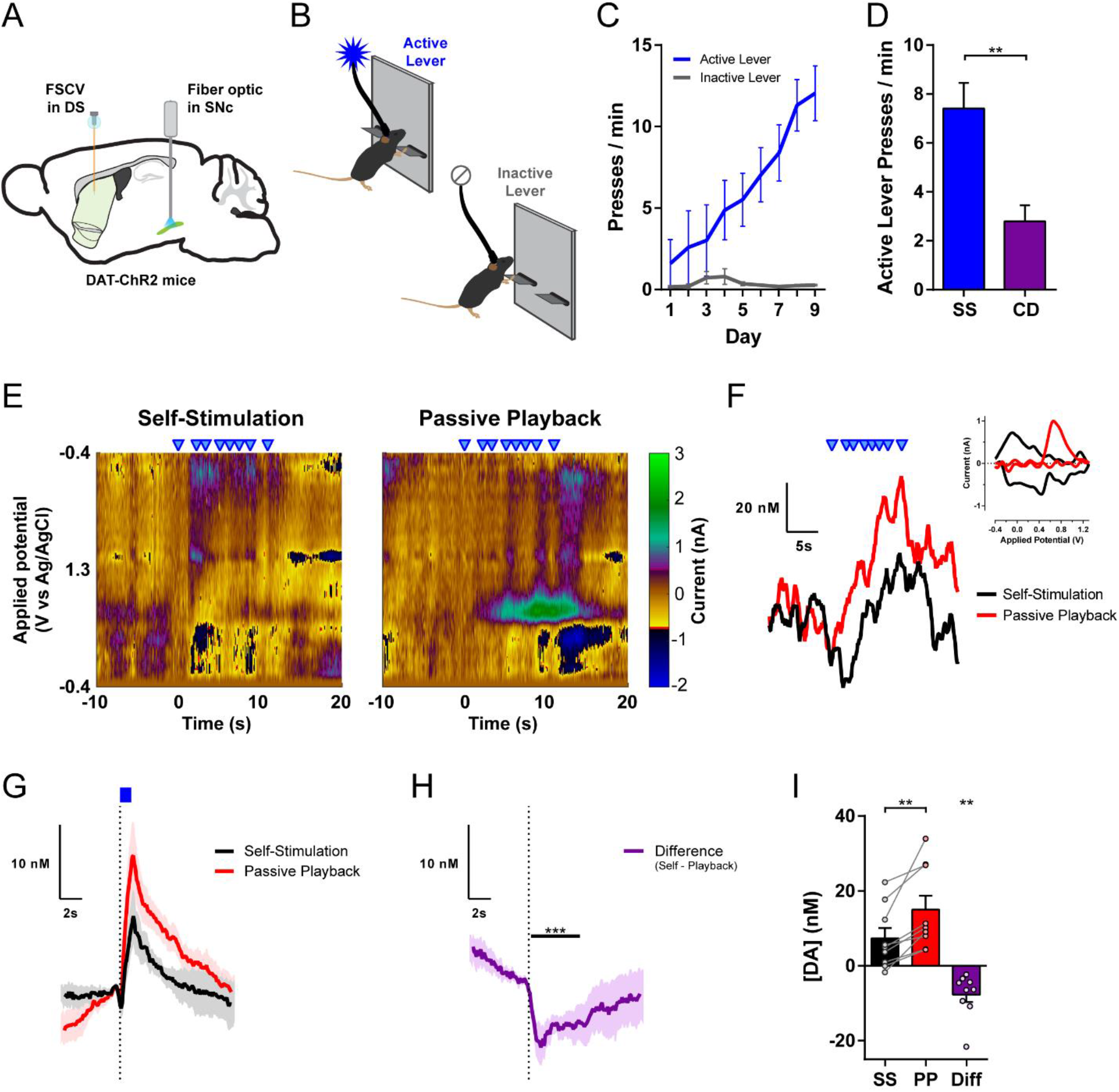
Goal-Directed Action Suppresses Nigrostriatal Dopamine Release During Opto-ICSS. (**A**) Schematic of experimental preparation for optogenetic stimulation of SNc dopamine neurons and FSCV recording in dorsal striatum. (**B**) Opto-ICSS behavioral task schematic. (**C**) Acquisition of opto-ICSS (*n* = 9 mice; two-way repeated-measures ANOVA: main effect of Lever, *F*_1,8_= 16.84, *P* = 0.0034; main effect of Day, *F*_8,64_= 9.314, *P* < 0.0001; Lever by Day interaction, *F*_8,64_ = 10.63, *P* < 0.0001; Active Lever significantly greater than Inactive Lever for Days 4-9, *P*s ≤ 0.008). (**D**) Contingency degradation test: 30 min of opto-ICSS followed by 30 min contingency degradation test phase (*n* = 6 mice; paired t test, *t*_5_= 5.441, *P* = 0.0028). (**E**) Representative voltammetric pseudocolor plots from a bout of stimulations (blue triangles) during the Self-Stimulation phase (left) and the matched stimulations from the Passive Playback phase (right). (**F**) Dopamine responses to the series of stimulations in each session phase from the example in (E). Inset: cyclic voltammograms. (**G**) Mean change in dopamine concentration evoked by Self-Stimulation and Passive Playback stimulations (*n* = 9 mice). (**H**) Difference trace: Self-Stimulation minus Passive Playback, from traces in (G). Black bar indicates post-stimulation time points with significant difference vs. 0 (permutation test, *P* = 0.0001). (**I**) Mean change in dopamine concentration (Self-Stimulation vs. Passive Playback paired t test, *t*_8_= 3.923, *P* = 0.0044; equivalently, Difference vs. 0 one-sample t test: *t*_8_= 3.923, *P* = 0.0044). SS, Self-Stimulation; CD, Contingency Degradation; PP, Passive Playback; Diff, Difference (Self minus Playback). All error bars are SEM, same for below unless stated otherwise. See also Figure 1—figure supplement 1.

To examine the extent to which this behavior is indeed goal-directed, a subset of mice underwent a contingency degradation test (Witten, et al., 2011; Koralek, et al., 2012; Clancy et al., 2014; Neely et al., 2018). During this test phase, stimulation was decoupled from the lever-pressing action and instead delivered non-contingently at a rate yoked to that animal’s own stimulation rate from the preceding self-stimulation phase (Methods). The mice significantly reduced their performance rate (Figure 1D, Figure 1—figure supplement 1G-H), indicating that they readily learned that their action was no longer required to earn stimulation. This demonstrates that nigrostriatal dopamine neuron self-stimulation under this simple fixed-ratio schedule of continuous reinforcement (CRF) is sensitive to changes in the action-outcome contingency, which is an established operational hallmark of goal-directed behavior (Yin et al., 2008; Balleine, 2019).

To investigate whether goal-directed action affects the nigrostriatal dopamine response to the consequence of that action, we used FSCV to record subsecond dopamine transmission in behaving mice during sessions that included two phases: In the Self-Stimulation phase, as in prior opto-ICSS training, mice earned optogenetic stimulation for each Active lever press (Figure 1B). In the subsequent Passive Playback phase, the levers were retracted, and the mice received non-contingent stimulations, with timestamps identical to the stimulations that each individual had self-administered in its Self-Stimulation phase. Thus, in this entirely within-subject design, we recorded at the same striatal location with the same chronically implanted electrode, with each animal yoked to its own performance, receiving the same temporal sequence of stimulations across both phases of the session, delivered to the same site within the SNc using identical optogenetic stimulation parameters to directly depolarize these nigrostriatal dopamine neurons (Figure 1A and 1E-F).

We observed a remarkably robust difference between the amplitude of Self-Stimulated dopamine release and the significantly greater amplitude evoked by the non-contingent Passive Playback stimulation (Figure 1F-I). All individual mice (9/9, 100%) exhibited less dopamine release when evoked as the consequence of their own action; this difference was significant at the individual level in 7 of 9 mice (*P*s < 0.0001) and was a trend in the same direction for the remaining 2 mice (*P*s = 0.0623 and 0.0825). Although the free-operant opto-ICSS task was designed to minimize discrete external cues, it nevertheless is possible that the offset of a previous stimulation essentially could serve as a stimulus that might elicit the next lever-pressing response. However, when we isolated the initiation of lever-pressing bouts using an inter-stimulation interval (ISI) criterion of at least 10 s since the previous stimulation, this subset of stimulations still showed a significant difference between Self-Stimulated and non-contingent Playback-evoked dopamine release (Figure 1—figure supplement 1I-K). This finding further indicates that optogenetically evoked dopamine release is lower when it is the outcome of self-initiated, goal-directed actions.

### Nigrostriatal Dopamine Signals Action-Outcome Prediction Errors

The reward prediction error theory implies decreased dopamine responses to expected versus unexpected outcomes (Houk et al., 1995; Montague et al., 1996; Schultz et al., 1997; Cohen et al., 2012; Eshel et al., 2016; Sutton and Barto, 2018). Nevertheless, the relative difference we observed does not alone resolve whether dopamine release is in fact inhibited by the animal’s action. To address this question, we recorded additional sessions in which a random 20% of Active lever presses did not yield stimulation, instead causing a 5-s timeout period during which no further stimulation could be earned (Figure 2A). During these Omission Probes, there was a clear dip in dopamine below baseline levels (Figure 2B-C), consistent with a neurochemical instantiation of a negative prediction error (Hart et al., 2014). Indeed, the timecourse for this Omission Probe dip was remarkably similar to the digital subtraction (“Difference Trace”) of the Self-Stimulated dopamine response minus the Passive Playback response (Figure 1H; overlaid in Figure 2D). This Omission Probe dip was not merely an artifact of FSCV background subtraction (Hamid et al., 2016), where reuptake during the stimulation-free timeout period might follow an elevated baseline from several preceding stimulations. Rather, a significant dip below baseline was still prominent for the subset of Omission Probes with a minimum latency of at least 5 s since the previous stimulation, whereas no such decrease was detected at the equivalent time points from the Playback phase (Figure 2—figure supplement 1A-B). Furthermore, additional lever presses during an ongoing stimulation augmented the suppression of Self-Stimulated dopamine release, and similarly, additional presses during an Omission Probe timeout period prolonged the duration of the dip below baseline (Figure 2—figure supplement 1C-F). *In vivo* extracellular electrophysiological recording further revealed reduced somatic firing in optogenetically-identified SNc dopamine neurons in response to action-evoked optogenetic Self-Stimulation relative to non-contingent Passive Playback stimulation (Figure 2— figure supplement 1G-L). Collectively, these results demonstrate that the action indeed causes inhibition of dopamine transmission.

**Figure 2.**
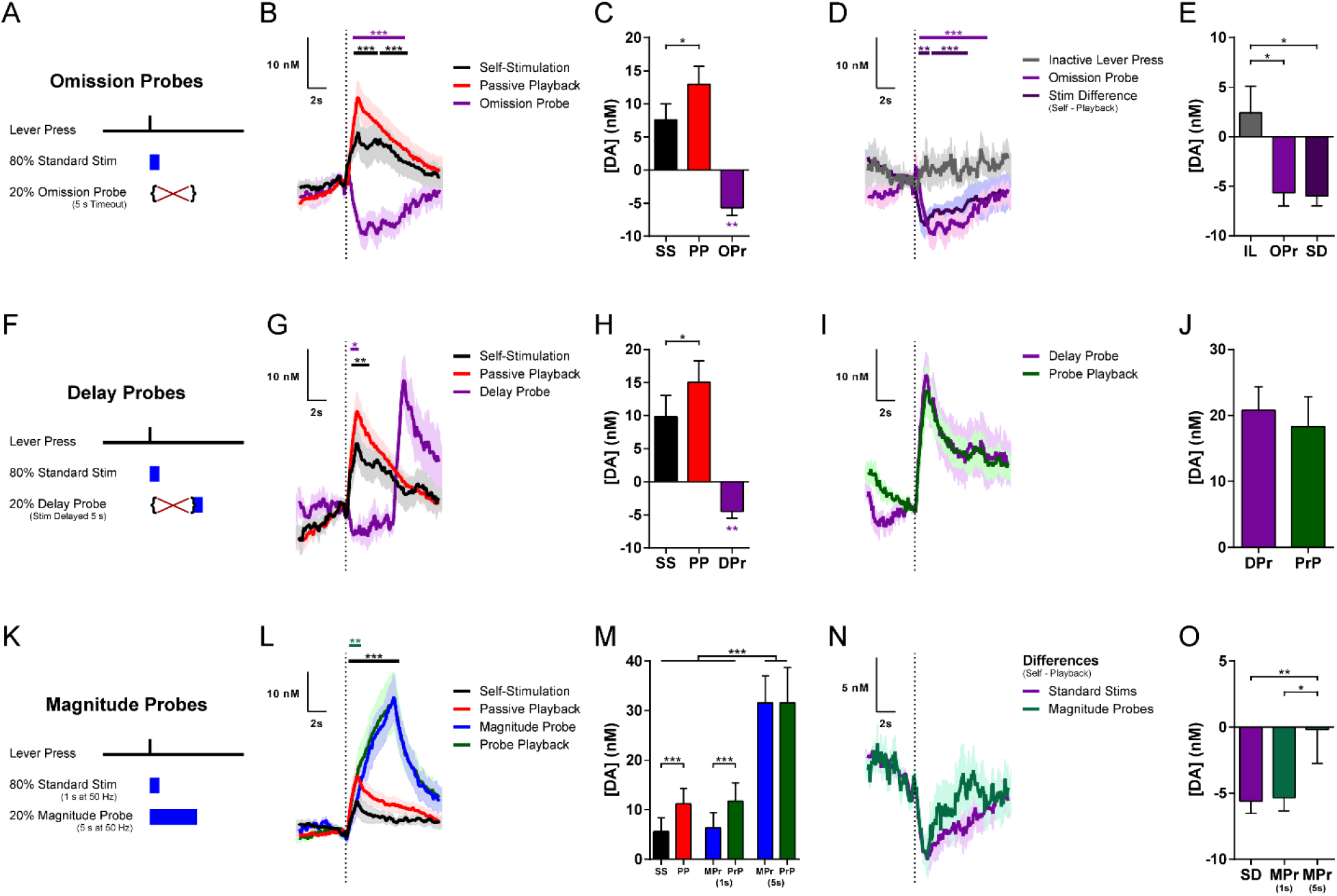
The Inhibition of Dopamine is Action-Specific and Temporally Precise. (**A**) Task schematic for Omission Probe sessions. (**B**) Mean change in dopamine concentration for stimulations and probes in Omission Probe sessions (*n* = 8 mice; permutation tests: Omission Probe vs. 0, magenta bar, *P* = 0.0001; Self-Stimulation vs. Playback, black bar, *P* = 0.0003 for first time cluster and *P* = 0.0001 for second cluster). (**C**) Mean change in dopamine concentration in Omission Probe sessions (Self-Stimulation vs. Passive Playback, paired t test, *t*_7_= 3.114, *P* = 0.0170; Omission Probe vs. 0, one-sample t test: *t*_7_= 4.810, *P* = 0.0019). (**D**) Mean change in dopamine concentration for Inactive Lever presses, overlaid on Omission Probes from (B) and Difference traces (Self Stimulation minus Playback) from Fig. 1H *(n* = 7 mice; permutation tests: Inactive press vs. Omission Probe, magenta bar, *P* = 0.0001; Inactive press vs. Stim Difference, purple bar, *P* = 0.007 for first time cluster and *P* = 0.0007 for second cluster. (**E**) Mean change in dopamine concentration for Inactive Lever presses versus Omission Probes from (C) and stimulation Differences from Fig. 1I (one-way repeated-measures ANOVA, *F*_2,12_= 7.419, *P* = 0.0080; Tukey’s multiple comparisons tests: Inactive Lever vs. Omission Probe, *P* = 0.0133; Inactive Lever vs. Stimulation Difference, *P* = 0.0174). (**F**) Task schematic for Delay Probe sessions. (**G**) Mean change in dopamine concentration for Standard Stimulations and probes in Delay Probe sessions (*n* = 8 mice; permutation tests: Delay Probe timeout period vs. 0, magenta bar, *P* = 0.0103; standard Self-Stimulation vs. Playback, black bar, *P* = 0.0001). (**H**) Mean change in dopamine concentration in Delay Probe sessions (Self-Stimulation vs. Passive Playback, paired t test, *t*_7_= 2.962, *P* = 0.0210; Delay Probe vs. 0, one-sample t test: *t*_7_= 4.341, *P* = 0.0034). (**I**) Mean change in dopamine concentration for Delay Probes and Probe Playback, each aligned to stimulation onset. (**J**) Mean change in dopamine concentration for Delay Probe stimulation and Probe Playback. (**K**) Task schematic for Magnitude Probe sessions. (**L**) Mean change in dopamine concentration for standard stimulations and probes in Magnitude Probe sessions (*n* = 8 mice; permutation tests: Magnitude Probe vs. Probe Playback, teal bar, *P* = 0.0052; standard Self-Stimulation vs. Playback, black bar, *P* = 0.0001). (**M**) Mean change in dopamine concentration in Magnitude Probe sessions (two-way repeated-measures ANOVA: main effect of Session Phase (Self vs. Playback), *F*_1,7_= 6.769, *P* = 0.0353; main effect of Stimulation Type (Standard, early and late Magnitude Probe), *F*_2,14_= 31.32, *P* < 0.0001; Session Phase by Stimulation Type interaction, *F*_2,14_= 7.724, *P* = 0.0055; Sidak’s multiple comparisons tests: late Magnitude Probes vs. Standard Stimulations and vs. early Magnitude Probes within both Self and Playback phases, all *P*s < 0.0001; Self vs. Playback for Standard Stimulations, *P* = 0.0006; Self vs. Playback for early Magnitude Probes, *P* = 0.0008). (**N**) Difference traces: Self-Stimulation minus Passive Playback for Standard Stimulations and Magnitude Probes, from traces in (L). (**O**) Mean Differences comparing session phases (Self-Stimulation minus Passive Playback) for Standard Stimulations and Magnitude Probes at early (1 s) and late (5 s) time points from Difference traces in (N). (One-way repeated-measures ANOVA, *F*_2,14_= 7.653, *P* = 0.0057; Tukey’s multiple comparisons tests: Standard Stimulation Differences vs. late Magnitude Probe Differences, *P* = 0.0099; early vs. late Magnitude Probe Differences, *P* = 0.0136). SS, Self-Stimulation; PP, Passive Playback; OPr, Omission Probe; IL, Inactive Lever; SD, Standard Stimulation Difference (Self minus Playback); DPr, Delay Probe; PrP, Probe Playback; MPr, Magnitude Probe. See also Figure 2— figure supplement 1.

It recently has been reported that some dopamine neurons transiently reduce their firing rate during certain types of spontaneous movement (Dodson et al., 2016; da Silva et al. 2018; Coddington and Dudman, 2018). We therefore considered the possibility that the action-induced suppression observed in our recordings may be a generalized inhibition following any lever pressing action, regardless of whether that action is associated with a particular reinforcing outcome. However, we found no such inhibition in the instances when the animal pressed the Inactive lever, which had never been reinforced throughout training (Figure 2D-E), indicating that the action-induced inhibition of dopamine release is specific to the typically reinforced action and conveys a bona fide prediction-error signal. Furthermore, because the sensory feedback from pressing the Active and Inactive levers is rather similar to the animals, these results confirmed that the dopamine inhibition results from the expectations associated with specific self-initiated, goal-directed action but not simply a conditioned sensory cue.

To examine the temporal specificity of this action-induced suppression, we recorded Delay Probe sessions in which 20% of Active lever presses instead resulted in stimulation that was delayed by 5 s (Figure 2F). The initial 5 s of this delay period was equivalent to the timeout period of the Omission Probes, and we again observed a dip in dopamine below baseline (Figure 2G-H). When the probe stimulation was finally delivered at the end of the delay period, there now was a high amplitude of dopamine release that did not differ from the corresponding Playback stimulations (Figure 2I-J). Because these Delay Probes were randomly interleaved throughout the Self-Stimulation phase, this indicates that there is not a global suppression of dopamine neuron excitability throughout the whole context of the Self-Stimulation phase. Rather, this action-induced inhibition is precisely timed to counteract the expected consequence of that action, namely the immediate stimulation that is its typical outcome.

We further determined the nature of this action-induced inhibition in Magnitude Probe sessions, where 20% of Active lever presses yielded 5 s of stimulation rather than the standard 1-s stimulation used throughout training (Figure 2K). These increased Magnitude Probes indeed evoked much greater dopamine release, as expected for longer-duration stimulation (Figure 2L-M). Closer examination of the time course of these dopamine responses also revealed a transient suppression during the self-stimulated Magnitude Probes that was restricted to the first second or so following stimulation onset, but no longer differed from Probe Playback by the end of the 5-s probe stimulation. This brief inhibition also was borne out by the transient dips and similar overall time courses in the Difference traces for both the Magnitude Probes and the standard 1-s stimulations, comparing each type of Self-Stimulation to their respective Playback stimulations (Figure 2N-O). This again highlights the timing and duration specificity of the action-induced suppression, and suggests that there is not a global inhibition of dopamine throughout the Self-Stimulation context. Together, these data suggest that nigrostriatal dopamine can encode a reward prediction error signal for individual goal-directed action and its expected outcome.

### Sequence-Specific Suppression of Nigrostriatal Dopamine Release

In real life, goals are seldom achieved by a single action but instead mostly through a series of actions organized in spatiotemporal sequences (Gallistel, 1980; Jin and Costa 2010; Jin et al., 2014; Geddes et al., 2018). Having established that the observed prediction error-like suppression of nigrostriatal dopamine is action-specific and temporally restricted, we next turned to the question of whether such regulation of dopamine transmission reflects hierarchical control over learned action sequences (Lashley, 1951; Gallistel, 1980; Geddes et al., 2018). To this end, we trained a separate cohort of mice to perform a spatiotemporally heterogeneous action sequence, pressing the Left and then Right lever (LR) to earn optogenetic nigrostriatal dopamine neuron self-stimulation (See Methods; Figure 3A, Figure 3—figure supplement 1A-B). As mice increased the number of stimulations earned across days of training (Figure 3B), their behavior exhibited several indications of successfully learning this LR action sequence: They increased both their probability of correctly completing a sequence by transitioning to a Right lever press following each Left lever press and their probability of reinitiating with a Left lever press following each stimulation (Figure 3C). Their duration to complete these LR sequences was shorter than the post-reinforcer reinitiation latency (Figure 3D), and the proportion of correct LR sequences increased relative to other non-reinforced press pairs (Figure 3E). The total presses per sequence and the number of consecutive presses on either lever both decreased throughout training, collectively contributing to an increase in overall efficiency (Figure 3—figure supplement 1C-F). Therefore, rather than simply associating the reinforcing outcome with the most proximal action at the Right lever, the animals’ behavior suggested that they indeed concatenated the distinct action elements into chunked action sequences. Furthermore, the mice significantly reduced their LR sequence performance during a contingency degradation test (Figure 3—figure supplement 1G), indicating that these chunked action sequences also were goal-directed.

**Figure 3.**
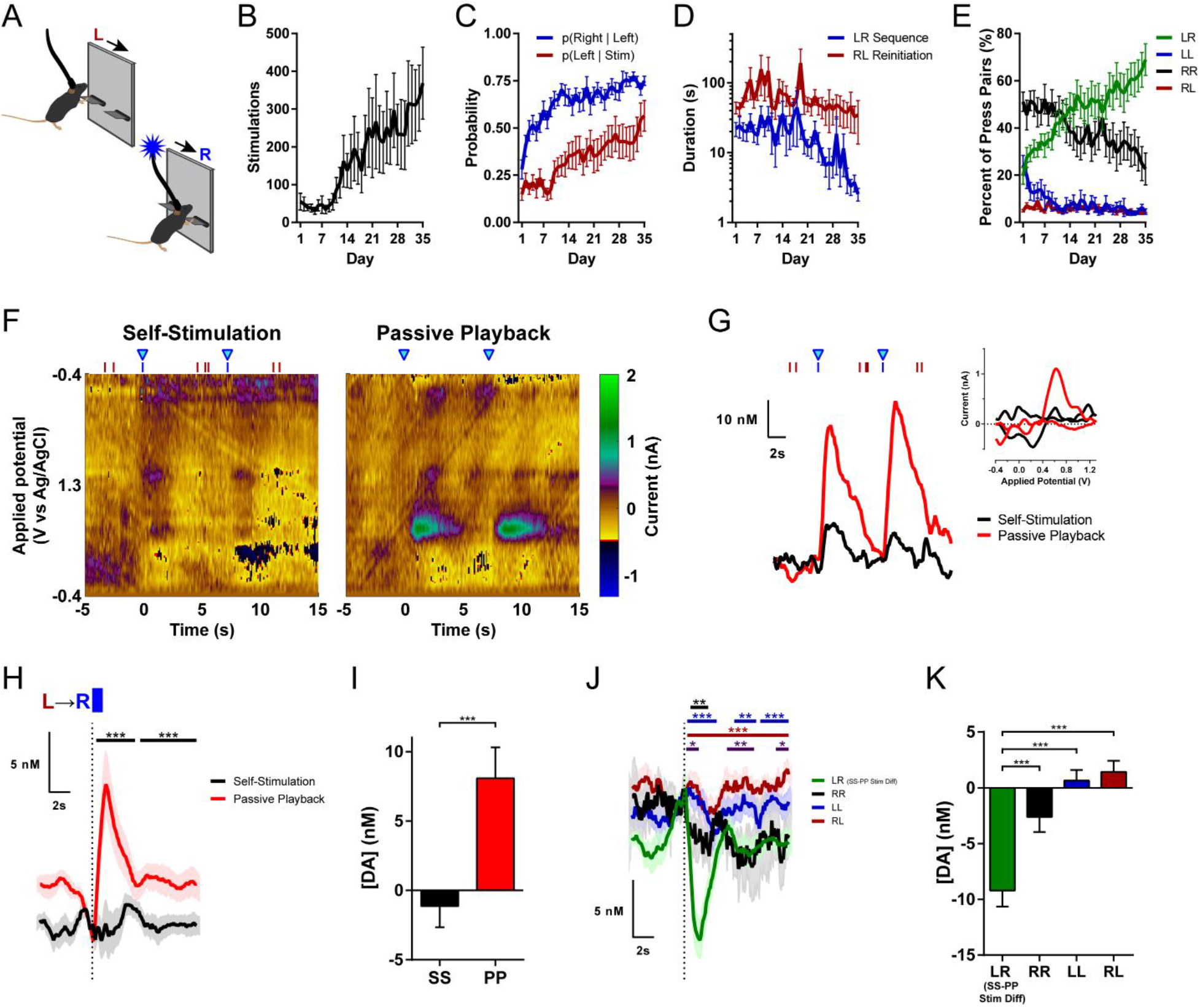
Sequence-Specific Inhibition of Nigrostriatal Dopamine during Performance of Learned LR Sequence. (**A**) Left-Right Sequence Self-Stimulation task schematic. (**B**) Stimulations earned across days of training (*n* = 13 mice; one-way repeated-measures ANOVA: *F*_34,408_= 3.550, *P* < 0.0001). (**C**) Transition probabilities: probability of pressing Right lever after each Left lever press, and probability of reinitiating Left press after stimulation (two-way repeated-measures ANOVA: main effect of Day, *F*_34,408_= 8.838, *P* < 0.0001; main effect of Transition Type, *F*_1,12_= 47.18, *P* < 0.0001; Dunnett’s multiple comparisons tests vs. Day 1: p(Right | Left) first significantly differs on Day 3, *P* = 0.0266; p(Left | Stim) first significantly differs on Day 14, *P* = 0.0265). (**D**) Median latencies to complete Left-Right sequences and to reinitiate a new sequence after previous stimulation (two-way repeated-measures ANOVA: main effect of Interval Type, *F*_1,12_= 18.79, *P* = 0.0010). (**E**) Relative frequency of each combination of lever press pairs (two-way repeated-measures ANOVA: main effect of Pair Type, *F*_3,36_= 32.58, *P* < 0.0001; Pair Type by Day interaction, *F*_102,1224_= 4.315, *P* < 0.0001). (**F**) Representative voltammetric pseudocolor plots during two Left-Right sequences for optogenetic intracranial Self-Stimulation (left) and corresponding stimulations from the Passive Playback phase (right). (**G**) Dopamine responses to the stimulations depicted in (F). Red and blue ticks denote left and right lever presses, respectively. Inset: cyclic voltammograms evoked by the first stimulation. (**H**) Mean dopamine concentration changes evoked by stimulation in each phase (*n* = 12 mice; permutation test, *P*s = 0.0001 for both time clusters). (**I**) Mean change in dopamine concentration (paired t test, *t*_11_= 6.403, *P* < 0.0001). (**J**) Mean dopamine concentration changes evoked by non-stimulated pairs of lever presses (5-s maximum inter-press interval within pair). Traces are aligned to the second press in each pair type. The LR sequence stimulation Difference (Self-Stimulation minus Passive Playback) is overlaid for comparison (green). (Permutation tests: LR Stim Difference vs. RR, black bar, *P* = 0.0046; LR Stim Difference vs. LL, blue bars, *P*s = 0.0002 for first time cluster, 0.0017 for second, and 0.0007 for third cluster; LR Stim Difference vs. RL, maroon bar, *P* = 0.0001; RR vs. RL, purple bars, *P*s = 0.0185 for first time cluster, 0.0021 for second, and 0.0163 for third cluster). (**K**) Mean change in dopamine concentration for each combination of non-reinforced press pairs during the Self-Stimulation phase and the LR sequence stimulation Difference (Self minus Playback). (One-way repeated-measures ANOVA, *F*_3,33_= 20.34, *P* < 0.0001; Tukey’s multiple comparisons tests: LR Stim Difference vs. RR, *P* = 0.0007; LR Stim Difference vs. LL and vs. RL, *P*s < 0.0001). Stim, Stimulation; LR, Left-Right sequence; RR, Right-Right; LL, Left-Left; RL, Right-Left; SS, Self-Stimulation; PP, Passive Playback; Diff, Difference. See also Figure 3—figure supplement 1.

We then recorded nigrostriatal dopamine transmission in these sequence-trained mice using the same within-subject manipulation comparing Self-Stimulation versus Passive Playback-evoked dopamine responses. We again found a robust suppression of the Self-Stimulated dopamine response (Figure 3F-I, Figure 3—figure supplement 1H-I), recapitulating the main result from the single-lever CRF cohort (Figure 1E-I). Importantly, no other combination of non-reinforced press pairs caused inhibition comparable to the difference between the Self-Stimulated versus Playback-evoked suppression of dopamine (Figure 3J-K), indicating that this inhibition was specific to the learned action sequence.

### Differential Regulation of Dopamine by Individual Actions within Learned Sequence

Beyond this sequence-type specificity, we further examined the question of whether dopamine transmission might reflect regulation at the level of individual action elements or instead at a higher sequence level in a hierarchy of behavioral control. For example, if regulated with each action element, we might expect similar inhibition for each individual Left and Right lever press, and summation of each to the full inhibition at outcome delivery. Alternatively, since animals chunked these action elements into fully concatenated action sequences, we might expect the action-induced inhibition of dopamine to begin at sequence initiation and persist throughout performance of this chunked action sequence. The results were inconsistent with either of these hypotheses, instead exhibiting a distinct form of sequence-specificity consistent with hierarchical control (Jin and Costa 2015; Geddes et al., 2018). Initiating Left lever presses did not cause any inhibition of dopamine, instead revealing a slight, albeit non-significant increase in dopamine release (Figure 4A-B). Similarly, additional recording sessions with probe stimulations delivered on 20% of initiating Left presses revealed no inhibition of dopamine evoked by these Left Probes (Figure 4C-H). Instead, these Left Probes actually evoked significantly greater dopamine release than their Playback (Figure 4F). At the individual animal level, 5 out of 8 mice (62.5%) showed a significant dopamine increase, and none showed a significant suppression. Outside of LR sequences, single Right lever presses (see Methods) did not result in inhibition of dopamine, in stark contrast with the full inhibition of Self-Stimulated versus Playback-evoked dopamine for correct LR sequences (Figure 4I-J). The dopamine response to Probe stimulations for these isolated Right presses also did not differ from their Playback, exhibiting no significant inhibition (Figure 4K-P). This same action at the Right lever therefore reveals highly distinct regulation of dopamine dynamics depending on the action’s membership within the learned sequence or not. These results indicate that the dopaminergic prediction errors are selective to the learned action sequence and reflect sequence-level hierarchical control over instrumental behavior.

**Figure 4.**
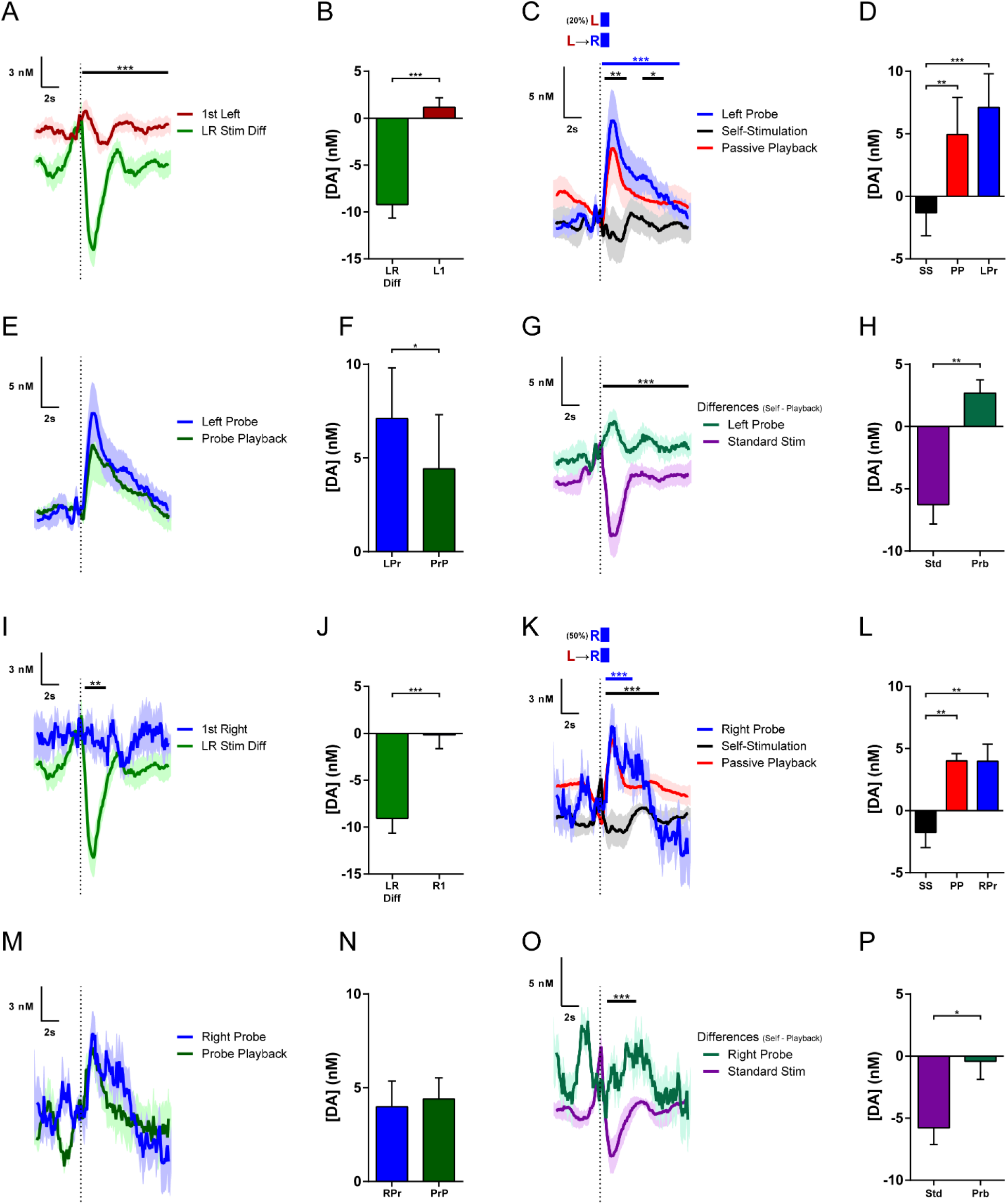
Different Regulation of Dopamine by Individual Actions of the LR Sequence. (**A**) Mean dopamine concentration change to first Left lever press following previous stimulation during Left-Right sequence task performance, overlaid with the Difference Trace (Self minus Playback) for LR sequence stimulations from Fig. 3J for comparison (*n* = 12 mice; permutation test, *P* = 0.0001). (**B**) Mean change in dopamine concentration for first Left lever presses and LR sequence stimulation Difference (*t*_11_= 6.325, *P* < 0.0001). (**C**) Mean dopamine concentration changes in Left-Right sequence sessions with Left Lever Probes (*n* = 8 mice; permutation tests: Left Probe vs. Standard LR Self-Stimulation, blue bar, *P* = 0.0001; Standard LR Self-Stimulation vs. Playback, black bars, *P*s = 0.0049 for first and 0.011 for second time clusters, respectively). (**D**) Mean change in dopamine concentration in Left Probe sessions (one-way repeated-measures ANOVA, *F*_2,14_= 17.19, *P* = 0.0002; Tukey’s multiple comparisons tests: Left Probe vs. LR Self-Stimulation, *P* = 0.0002; LR Self-Stimulation vs. Playback, *P* = 0.0024). (**E**) Mean dopamine concentration changes in Left Probes and the Probe Playback. (**F**) Mean change in dopamine concentration to Left Probes and Probe Playback (paired t test, *t*_7_= 2.519, *P* = 0.0399). (**G**) Difference traces from Left Probe Sessions: Self-Stimulation minus Passive Playback for Standard LR Stimulations and Left Probes, from traces in (C) and (E); (permutation test, *P* = 0.0001). (**H**) Mean Differences comparing session phases (Self-Stimulation minus Passive Playback) for Standard LR Stimulations and Left Probes (paired t test, *t*_7_= 5.125, *P* = 0.0014). (**I**) Mean dopamine concentration change to first Right lever press following previous stimulations, overlaid with the Difference Trace (Self minus Playback) for LR sequence stimulations from Fig. 3J for comparison. First Right press was an additional press on the Right lever after a previous stimulation, without approaching the Left lever (*n* = 11 mice; permutation test, *P* = 0.0012). (**J**) Mean change in dopamine concentration for first Right lever presses and LR sequence stimulation Difference (*t*_10_= 5.690, *P* = 0.0002). (**K**) Mean dopamine concentration changes in Left-Right sequence sessions with Right Lever Probes where animal did not approach the Left lever in the preceding inter-stimulation interval (*n* = 10 mice; permutation tests: Right Probe vs. Standard LR Self-Stimulation, blue bar, *P* = 0.0001; Standard LR Self-Stimulation vs. Playback, black bar, *P* = 0.0001). (**L**) Mean change in dopamine concentration in Right Probe sessions (one-way repeated-measures ANOVA, *F*_2,18_= 10.47, *P* = 0.0010; Tukey’s multiple comparisons tests: Right Probe vs. LR Self-Stimulation, *P* = 0.0026; LR Self-Stimulation vs. Playback, *P* = 0.0024). (**M**) Mean dopamine concentration changes in Right Probes and the Probe Playback. (**N**) Mean change in dopamine concentration to Right Probes and Probe Playback. (**O**) Difference traces from Right Probe Sessions: Self-Stimulation minus Passive Playback for Standard LR Stimulations and Right Probes, from traces in (K) and (M); (permutation test, *P* = 0.0002). (**P**) Mean Differences comparing session phases (Self-Stimulation minus Passive Playback) for Standard LR Stimulations and Right Probes (paired t test, *t*_9_= 3.080, *P* = 0.0131). LR Stim Diff, Left-Right Stimulation Difference (Self minus Playback); L1, 1^st^ Left Press; R1, 1^st^ Right Press; SS, Self-Stimulation; PP, Passive Playback; LPr, Left Probe; PrP, Probe Playback; Std, Standard LR Stimulation; Prb, Probe; RPr, Right Probe.

## DISCUSSION

Overall, we have demonstrated that nigrostriatal dopamine transmission to reinforcing outcomes is strongly suppressed when this outcome is the expected consequence of the animal’s own action. This inhibition of outcome-evoked dopamine following self-initiated actions parallels commonly observed reward prediction errors in explicit stimulus-outcome and stimulus-response behavioral contexts. The current results therefore expand this phenomenon to include action-outcome prediction errors that support instrumental associations underlying self-initiated goal-directed behavior. This action-outcome prediction error was specific to the typically reinforced action, temporally restricted to counteract the expected consequence of that action, and exhibited sequence selectivity consistent with a high level of hierarchical control over chunked action sequences. The prediction errors signaled by dopamine transmission therefore reflect not only expectations associated with Pavlovian cues or behavioral responses to such discrete stimuli, but also the expected outcomes of self-initiated instrumental actions and sequences. Compelling behavioral and neural evidence for action chunking also is well established (Graybiel, 1998; Hikosaka, 1998; Matsumoto et al., 1999; Barnes et al., 2005; Jin and Costa, 2010; Wassum et al., 2012; Jin et al., 2014; Jin and Costa 2015; Collins et al., 2016; Geddes et al., 2018), but the mechanisms subserving such sequence learning remain poorly understood. While the exact role of nigrostriatal dopamine throughout sequence acquisition requires further direct investigation, the current results demonstrate that the performance of well-learned action sequences entails distinct dopamine dynamics for actions within these sequences. That nigrostriatal dopamine transmits specific action-outcome prediction errors and exhibits sequence-dependent hierarchical regulation provides critical new insight into these important neuromodulatory dynamics in goal-directed behavioral control, an under-examined domain of instrumental action beyond spontaneous movement of unknown purpose and responding to reward-predictive cues.

Several aspects of our opto-ICSS experimental design conferred distinct advantages for examining the regulation of nigrostriatal dopamine dynamics in goal-directed behavior. The current study used an entirely within-subject design and direct optogenetic excitation to selectively stimulate dopamine neurons and record dopamine transmission at identical locations within a given animal, in contrast to previous ICSS studies that used non-selective electrical stimulation of the midbrain and compared dopamine release between trained versus naïve animals (Garris et al., 1999; Kilpatrick et al., 2000) or did not include temporally matched non-contingent playback (Owesson-White et al., 2008; Rodeberg et al., 2016; Covey and Cheer, 2019). Traditional procedures with natural reward invariably require additional consummatory actions such as magazine approach or licking for reward retrieval, which itself might regulate dopamine dynamics and complicate the data analyses. Although selective optogenetic stimulation lacks the specific sensory features such as flavor that typically define the identity of natural reward outcomes (Kruse et al., 1983; Corbit and Janak, 2007; Collins et al., 2016; Takahashi et al., 2017; Keiflin et al., 2018; Balleine, 2019), the direct intracranial delivery permitted precise temporal control over outcome receipt. Furthermore, direct optogenetic stimulation also bypasses afferent circuitry representing any natural reward itself, permitting the current focus on regulation of dopamine by specific action-associated expectancies. These features of the current design collectively yielded results consistent with nigrostriatal dopamine transmitting an action-outcome prediction error signal.

Although direct optogenetic stimulation indeed approaches an essentially identity-less outcome (Wise, 2002), this outcome delivery does coincide with sensory feedback during the action, such as somatosensory contact or auditory feedback from pressing the lever. However, these sensory reafferents are comparable for inactive lever presses or other non-reinforced action sequences, and therefore cannot account for the selective suppression of dopamine evoked as the consequence of reinforced actions (Figures 2D-E and 3J-K). Indeed, the distinct regulation of dopamine to the same action depending on sequence membership (Figure 4I-P) again provides clear evidence that the observed suppression was due to specific action expectancies rather than sensory feedback. Whereas the suppression of outcome-evoked dopamine release is therefore unlikely accounted for by different sensory features between the session phases, this action-induced suppression may instead share important commonalities with efference copy (or corollary discharge) phenomena widely observed in numerous other sensorimotor systems throughout the nervous systems of many different species (Wolpert et al., 1995; Crapse and Sommer, 2008; Schneider et al., 2018; Wurtz, 2018). Indeed, the current results provide evidence that a learned, sequence-level efference copy can suppress the neurochemical consequence of the complete action sequence, distinct from the regulation by individual action elements. These findings align with the recent demonstration of dopaminergic prediction errors for evaluating sequential sensorimotor control relative to internal performance templates (Gadagkar et al., 2016), and are broadly consistent with the prominent role proposed for efference copies in striatal-dependent learning (Redgrave and Gurney, 2006; Fee, 2014).

The current study’s recordings targeted the dorsomedial striatum (DMS), which is widely implicated in goal-directed instrumental behavior (Yin et al., 2005; Yin et al., 2008; Gremel & Costa, 2013; Balleine, 2019; Matamales et al., 2020). A natural next question is whether regulation of dopamine dynamics differs in other striatal subregions. Recent work found an attenuation of mesolimbic dopamine release in the nucleus accumbens core during self-paced opto-ICSS of ventral tegmental area dopamine neurons, albeit without comparison to temporally matched non-contingent playback stimulation (Covey & Cheer, 2019). Together, this finding and the present study extend earlier work reporting suppression of both mesolimbic (Garris et al., 1999) and nigrostriatal dopamine (Kilpatrick et al., 2000) evoked by non-selective electrical self-stimulation in trained animals versus non-contingent playback in naïve animals. Further, in a discrete-trial, cued task variant, Covey and Cheer (2019) also found an attenuation of optogenetically stimulated dopamine release and a concomitant increase in cue-evoked release, consistent with classic reward prediction errors in natural reward contexts (Schultz et al., 1997). Indeed, another recent study found predominant prediction-error responses in dopamine axonal activity throughout much of the ventral, dorsomedial, and dorsolateral striatum (DLS) in a cued discrimination task for water reward (Tsutsui-Kimura et al., 2020). Here, a notable difference in the DLS was a lack of dips below baseline despite similarly suppressive effects of reward expectation across regions. Based on these collective findings, we therefore would predict that most effects observed within the DMS in the current study would be largely similar in the accumbens core (Covey & Cheer, 2019), and although we also would expect suppression in the DLS, we also might not expect negative prediction errors to cause dips below baseline there (Tsutsui-Kimura et al., 2020). In contrast, the predictions are perhaps less clear for aspects of the accumbens shell and the caudal-most tail of the striatum, where distinct and surprising dopamine dynamics have been revealed particularly in aversive domains (de Jong et al., 2018; Menegas et al., 2018; Steinberg et al., 2020). Overall, potential heterogeneity of dopamine signaling across striatal subregions remains an important topic of investigation.

Uncovering the circuit mechanisms responsible for this dopaminergic action-outcome prediction error also remains an important open question for future research. The current results constrain candidate mechanisms to those with fairly rapid onset, transient duration, and sufficiently strong inhibition to suppress or shunt even direct optogenetic depolarization. Nigrostriatal dopamine neurons receive monosynaptic inputs from all basal ganglia nuclei (Watabe-Uchida et al., 2012; Lerner et al., 2015; Menegas et al., 2015), the majority of which are predominantly inhibitory GABAergic projections (Tepper and Lee, 2007; Brazhnik et al., 2008; Evans et al., 2020). Striatal, pallidal, and nigral basal ganglia nuclei contain many cells exhibiting prominent activity related to action sequence initiation, termination, and transitions (Jin and Costa, 2010; Jin et al., 2014; Geddes et al., 2018), as well as action-outcome value information (Samejima et al., 2005; Lau and Glimcher 2008; Hong and Hikosaka, 2008; Roesch et al., 2009; Tachibana and Hikosaka, 2012; Kim et al., 2017) that may converge and contribute to these dopamine neuron computations. Recent studies have suggested that dynamic nigrostriatal dopamine might regulate ongoing actions (Jin and Costa, 2010; Barter et al. 2015; Panigrahi et al. 2015; da Silva et al. 2018) and bias online action selection (Howard et al. 2017). The current results revealed that nigrostriatal dopamine can encode action-outcome prediction errors critical for action learning. Together they underscore the importance of dopamine for action selection at short as well as long timescales, and have important implications in many neurological disorders such as Parkinson’s disease, schizophrenia, and addiction.

## MATERIALS AND METHODS

| Key Resources Table |  |  |  |  |
| --- | --- | --- | --- | --- |
| Reagent type (species) or resource | Designation | Source or reference | Identifiers | Additional information |
| genetic reagent (*Mus musculus*) | DAT-cre (*Slc6a3*) | Jackson Laboratory | RRID:IMSR_JAX:020080 |  |
| genetic reagent (*Mus musculus*) | Ai32 (RCL-ChR2(H134R)/eYFP) | Jackson Laboratory | RRID:IMSR_JAX:024109 |  |
| strain, strain background (Adeno-associated virus) | AAV5-EF1a-DIO-ChR2(H134R)-mCherry | UNC vector core | RRID:SCR_002448 |  |
| antibody | anti-tyrosine hydroxylase (rabbit polyclonal) | Abcam | Cat#ab112; RRID:AB_297840 | (1:1000) |
| antibody | anti-GFP (chicken polyclonal) | Novus Biologicals | Cat#NB100-1614; RRID:AB_10001164 | (1:1000) |
| antibody | anti-rabbit Cy3 (donkey polyclonal) | Jackson ImmunoResearch | Cat#711-165-152; RRID:AB_2307443 | (1:250) |
| antibody | anti-chicken Alexa Fluor 488 (donkey polyclonal) | Jackson ImmunoResearch | Cat#703-545-155; RRID:AB_2340375 | (1:250) |

### Animals

All procedures were approved by the Institutional Animal Care and Use Committee at the Salk Institute for Biological Studies and were conducted in accordance with the National Institute of Health’s Guide for the Care and Use of Laboratory Animals. Experiments were performed using male and female mice, at least two months old, group-housed (2-5 mice / cage) on a 12 hr light/dark cycle (lights on at 6:00 am). *DAT-cre* mice (Jackson Laboratory # 020080; Zhuang et al., 2005) were either crossed with the Ai32 line (*RCL-ChR2(H134R)-EYFP*, Jackson Laboratory # 024109; Madisen et al., 2012) or injected with cre-dependent AAV in the SNc to selectively express channelrhodopsin-2 in their dopamine neurons.

### Surgical Procedures

Mice were anesthetized with isoflurane (3% induction, 0.5-1.5% sustained), their head shaved, and they were placed in a stereotaxic frame. The scalp was swabbed with 70% isopropyl alcohol and a povidine-iodine solution, and given a subcutaneous injection of bupivicaine (2 mg/kg) for local anesthesia. After a midline incision and leveling the skull, skulls were dried and coated with OptiBond adhesive and/or implanted with skull screws. Craniotomies were drilled over the dorsal striatum (+ 0.5-0.8 mm AP, 1.5 mm ML from bregma) for the voltammetric working electrode, the substantia nigra pars compacta (SNc:-3.1-3.3 mm AP, 1.3 mm ML) for the fiber optic(s), and an arbitrary distal site for the Ag/AgCl reference electrode. For *DAT-cre* mice not already crossed with the *Ai32* line, 300 nl of AAV5-EF1a-DIO-ChR2(H134R)-mCherry (UNC vector core) was injected into the SNc (4.1 mm ventral from dura; 100 nl/min), and the injection needle was left in place for 5 min before being slowing withdrawn (Howard et al., 2017). For all FSCV mice, the Ag/AgCl reference was inserted under the skull and cemented in place, and a carbon-fiber microelectrode (Clark, Sandberg et al., 2010) was lowered into the striatum (2.3-2.5 mm DV from dura) while applying a voltammetric waveform (see FSCV section) at 60 Hz for 10-15 min, and then at 10 Hz until the background had stabilized. A fiber optic (200 μm core; Sparta et al., 2011; Howard et al. 2017) was lowered targeting the ipsilateral SNc (3.8-4.1 mm DV). *DAT-cre x Ai32* mice received 1-s 50-Hz optical stimulation while striatal dopamine was recorded with FSCV to ensure electrode functionality and fiber placement. Mice subsequently trained in the Left-Right sequence task cohort (see Behavioral Training) also were implanted with a fiber optic over the contralateral SNc for bilateral stimulation. All implants were cemented to the skull along with a connector from the reference and working electrodes for later attachment to the FSCV head-mounted amplifier (headstage). For electrophysiological identification of dopamine neurons, *DATcre x Ai32* mice were implanted unilaterally in the SNc with an electrode array (Innovative Neurophysiology) with 16 tungsten contacts (2 x 8), 35 μm in diameter, spaced 150 μm apart within rows and 200 μm apart between rows. The array had a fiber optic directly attached, positioned ~300 μm from the electrode tips, to permit coupling to the laser for stimulation delivery (Jin and Costa 2010; Howard et al. 2017). The silver grounding wire was attached to a skull screw, and the array was affixed with dental cement. Mice received buprenorphine (1 mg/kg, s.c.) for analgesia and dexamethasone (2.5 mg/kg, s.c.) or ibuprofen in their drinking water for post-operative anti-inflammatory treatment, recovered in a clean home cage on a heating pad, were monitored daily for at least 3 days, and allowed to recover for at least 10 days before beginning behavioral training.

### Behavioral Training

Behavioral training was conducted in standard operant chambers (Med Associates) inside sound attenuating box, as previously described (Howard et al., 2017; Geddes et al., 2018). Mice were connected to the fiber optic patch cable from the laser (LaserGlow; 473 nm, ~5 mW measured before each session) and placed in the operant chamber, and optogenetic intracranial self-stimulation (opto-ICSS) sessions began with the insertion of two levers and the onset of a central house light on the opposite wall. The levers remained extended and the house light remained on for the duration of the 60 min sessions.

#### Continuous reinforcement cohort

Each press on the designated Active lever resulted in 1 s of optical stimulation (50 Hz, 10 ms pulse width) on a continuous reinforcement (CRF) schedule, other than additional presses during an ongoing stimulation train, which were recorded but had no consequence. Presses on the other, Inactive lever also were recorded but had no consequence. The sides of the Active and Inactive levers were counterbalanced relative to both the operant chamber and implanted hemisphere across mice, and remained fixed across training days for a given animal. Once mice reliably made at least 100 Active lever presses per session for 3 consecutive days, they also were connected to a voltammetry headstage before each session to allow habituation to behaving with this additional tethering. If a mouse failed to interact with the levers during its first 3 days of training, it was placed on food restriction overnight and a sucrose pellet was placed on the lever during its next behavioral session to encourage exploration. Once mice were reliably pressing the Active lever, they remained on *ad libitum* access to food and water in their home cages for all subsequent behavioral training and FSCV recordings. Mice were trained for at least 3 days while tethered to the FSCV headstage and meeting the behavioral criteria of at least 100 Active lever presses before FSCV recordings commenced (mean ± SEM = 11.1 ± 1.2 training days).

#### Left-Right sequence cohort

Mice in the Left-Right (LR) sequence cohort were initially trained on single-lever CRF opto-ICSS. For this cohort’s CRF training, only one lever was extended in each of two 30-min blocks per session (order counterbalanced across mice), and presses in both left- and right-lever blocks yielded the same 1-s, 50-Hz stimulation. To expedite this initial training stage, all mice in this cohort were food restricted prior to their first session, and were maintained at 85% their free-feeding baseline weight with ~2.5 g of standard lab chow per mouse in their home cage after the daily training sessions. Once mice made at least 100 presses in each block for 3 consecutive days, they were returned to *ad libitum* food access in their home cage, CRF training continued until they again met this 100-press criterion for another 3 days, and they then began training on the LR sequence task.

In LR sequence session, both levers were inserted at the start of the session and remained extended for the duration of the 60 min sessions. To receive stimulation (1 s, 50 Hz), mice now had to press the Left and then Right lever. No other combination of lever press pairs (Left-Left, Right-Right, or Right-Left) was reinforced with stimulation. After reaching the behavioral criterion of receiving at least 100 stimulations per session for 3 consecutive days, mice were habituated to tethering with the FSCV headstage, and received further training while tethered until they again met this 100-stimulation 3-day criterion and FSCV recording sessions commenced (mean ± SEM = 56.9 ± 7.6 training days). A subset of animals was trained under the same procedures to instead perform the Right-Left sequence as a spatial control, but we refer to the LR sequence throughout for simplicity. The hemisphere of the implanted FSCV recording electrode also was counterbalanced relative to this sequence direction across mice.

#### Contingency degradation

A contingency degradation test session began with 30 min of standard opto-ICSS (CRF for the CRF cohort, LR sequence task for the LR cohort). In the subsequent 30-min contingency degradation test phase, the levers remained extended, but stimulation was decoupled from task performance and instead was delivered regardless of whether the mice pressed any levers (Witten, Steinberg, et al., 2011; Koralek, Jin, et al., 2012; Clancy et al., 2014; Neely et al., 2018). For each mouse, the timing of these non-contingent stimulations during the test phase was matched to the time stamps of stimulations earned during that animal’s preceding opto-ICSS phase in the first half of the session, ensuring that the stimulation rate and distribution of inter-stimulation intervals were yoked within-subject to a given animal’s own opto-ICSS performance.

### Fast-Scan Cyclic Voltammetry (FSCV)

Striatal dopamine was recorded with *in vivo* FSCV in behaving animals as previously described (Clark et al., 2010; Hollon et al. 2014; Howard et al., 2017). Briefly, voltammetric waveform application consisted of holding the potential at the carbon-fiber electrode at-0.4 V relative to the Ag/AgCl reference between scans, and ramping to +1.3 V and then back to-0.4V at 400 V/s for each scan. Prior to the initial FSCV recording during opto-ICSS performance, this voltammetric waveform was applied at 60 Hz for at least one hour while mice were in a ‘cycling chamber’ outside the operant box, then at 10 Hz until the background current had stabilized. Mice then received experimenter-delivered optical stimulations (1 s, 50 Hz) to ensure electrode functionality.

For opto-ICSS sessions with FSCV recordings, electrodes were first cycled at 60 Hz for ~40 min and then at 10 Hz for at least 20 min until background current equilibration and throughout the opto-ICSS behavioral session. Mice received a series of 3 experimenter-delivered stimulations before and after the session to validate electrode functionality the day of each recording and for generating voltammetric training sets (see Statistical Analyses). The opto-ICSS session began at least 5 min after the final pre-session stimulation. The first half of each FSCV session consisted of a standard opto-ICSS phase (CRF for the CRF cohort, LR sequence task for the LR cohort) that was identical to the previous behavioral training sessions. At the conclusion of this active Self-Stimulation phase, the both levers retracted and the house light turned off for a 5 min interim period, followed by a Passive Playback phase in which mice received non-contingent stimulations with the same timing and stimulation parameters (1 s at 50 Hz) as in the active Self-Stimulation phase. The timing of these non-contingent Passive Playback stimulations was matched to the time stamps of stimulations earned during a given animal’s preceding Self-Stimulation phase, again ensuring that the stimulation rate and distribution of inter-stimulation intervals were identical across both the active and passive phases for a given animal. Mice in the LR sequence cohort also performed another 30 min of active LR sequence opto-ICSS following the Passive Playback phase to permit assessment of possible temporal order effects (Fig S3H-I).

Mice also underwent additional FSCV recordings in several types of probe sessions, including Omission, Delay, and Magnitude Probes for the CRF cohort, and Left and Right Lever Probes for the LR cohort. These FSCV sessions consisted of the same basic protocol described above, with active Self-Stimulation and non-contingent Passive Playback yoked within-subject. In Omission Probe sessions, 20% of presses on the typically Active lever did not yield stimulation, and instead caused a 5-s timeout period during which no further stimulation could be earned. This timeout period was not explicitly cued with any overt stimulus, other than the absence of the typical stimulation delivery. In Delay Probe sessions, 20% of presses on the Active lever resulted in stimulation that was delayed by 5 s. As for the Omission Probe timeout period, no further stimulation could be earned during this delay period. In Magnitude Probe sessions, 20% of Active lever presses yielded an increased magnitude of stimulation (5 s at 50 Hz). For the LR sequence cohort, the single-press probe sessions consisted of probe stimulations delivered on a random subset of first lever presses after previous reinforcement, in addition to continuous reinforcement for LR sequences as usual. For the Left Probe session, the next left lever press following the last reinforcement was stimulated with 20% probability. Due to the lower probability of an additional right lever press following a reinforcement, a right lever press following the last reinforcement was stimulated with 50% probability to collect enough probes for data analyses in the Right Probe session. Probe sessions were recorded at least 2 days apart, with standard opto-ICSS behavioral training sessions performed on the intervening days to allow return to baseline performance.

### In Vivo Electrophysiology

SNc dopamine neurons were recorded and identified as previously described (Jin & Costa, 2010; Howard et al., 2017). Briefly, neural activity was recorded using the MAP system (Plexon), and spike activities first were sorted online with a build-in algorithm. Only spikes with stereotypical waveforms distinguishable from noise and high signal-to-noise ratio were saved for further analysis. Behavioral training and recording sessions were conducted as described above for the CRF cohort. After recording the opto-ICSS session with active Self-Stimulation and Passive Playback phases, the recorded spikes were further isolated into individual units using offline sorting software (Offline Sorter, Plexon). Each individual unit displayed a clear refractory period in the inter-spike interval histogram, with no spikes during the refractory period (larger than 1.3ms). To identify laser-evoked responses, neuronal firing was aligned to stimulation onset and averaged across stimulations in 1-ms bins, and baseline was defined by averaging neuronal firing in the 1 s preceding stimulation onset. The latency to respond to stimulation was defined as the as the time to significant firing rate increase, with a threshold defined as > 99% of baseline activity (3 standard deviations). Only units with short response latency (< 10 ms) from stimulation onset and high correlation between spontaneous and laser-evoked spike waveforms (r > 0.95) were considered cre-positive, optogenetically identified dopamine neurons (Jin & Costa, 2010; Howard et al., 2017).

### Histology

Mice were anesthetized with ketamine (100 mg/kg, i.p.) and xylazine (10 mg/kg, i.p.), and the FSCV recording site was marked by passing a 70 μA current through the electrode for 20 s. Mice were transcardially perfused with 0.01 M phosphate-buffered saline (PBS) and then 4% paraformaldehyde (PFA) in PBS. Brains were removed, post-fixed in PFA at 4° for 24 hr, and then stored at 4° in a solution of 30% sucrose in 0.1 M phosphate buffer until ready for cryosectioning. Tissue was sectioned at 50 μm thickness on a freezing microtome, and striatal and SNc sections were mounted onto glass slides and coverslipped with AquaPoly mounting media containing DAPI (1:1000). Some sections also were processed for immunohistochemistry as previously described (Smith, Klug, Ross, et al., 2016; Geddes et al., 2018). Briefly, sections were washed 3 times for 15 min each in tris-buffered saline (TBS), and incubated for 1 hr in blocking solution containing 3% normal horse serum and 0.25% Triton-X 100 in TBS. Tissue was incubated for 48 hr in primary antibody against tyrosine hydroxylase (anti-TH, raised in rabbit, 1:1000, Abcam) and green fluorescent protein (anti-GFP, raised in chicken, 1:1000, Novus Biologicals) in this blocking solution at 4°, washed twice for 15 min in TBS and then for 30 min in the blocking solution, and then incubated for 3 hr in secondary antibody (anti-Chicken AlexaFluor 488 and anti-Rabbit Cy3, each 1:250, Jackson ImmunoResearch) in blocking solution. Finally, sections were washed 3 times for 15 min in TBS, mounted onto slides, and coverslipped with DAPI mounting media as above. Sections were imaged on a Zeiss LSM 710 confocal microscope with 10x and 20x objectives. All included FSCV animals were confirmed to have electrode placement in the dorsal striatum and fiber optics targeting the SNc.

### Statistical Analyses

FSCV data were low-pass filtered at 2 kHz, aligned to each lever press and/or stimulation onset, and background-subtracted using the mean voltammetric current in the 1 s prior to each aligned event of interest. Dopamine responses were isolated using chemometric principal component analysis with training sets consisting of cyclic voltammograms for dopamine, pH, and electrode drift (Keithley et al., 2009; Keithley & Wightman, 2011; Howard et al., 2017). Electrode-specific training sets were used for each animal and represented additional inclusion criteria for a given electrode, but similar results were obtained when reanalyzing data with a standardized training set across animals (Rodeberg et al., 2017). Changes in dopamine concentration were estimated based on average post-implantation electrode sensitivity (Clark, Sandberg, et al., 2010). No formal power analysis was conducted prior to experiments, but sample sizes were comparable to previous publications.

Mean changes in dopamine concentration summarized in bar graphs throughout the results analyzed time periods spanning 0.5-1.5 s following the aligned event onset. Analysis of the Magnitude Probes also included a late time point at 4.5-5.5 s after Probe onset, as did supplementary analysis of Omission Probes with versus without additional presses during the timeout period. For the LR sequence cohort, analysis of non-reinforced press pairs was restricted to pairs with short inter-press intervals (IPI < 5 s), consistent with the short duration of most LR sequences. Analysis of the non-reinforced single Left and Right lever presses was restricted to the first press following previous reinforcement, to match the press that could receive probe stimulation in the corresponding single-press probe sessions. Analysis of non-reinforced Right lever presses and Right Probe stimulations was restricted to those where the animal did not first approach the Left lever, as determined by examination of the video, to ensure that the right presses analyzed were individual actions and not part of a LR sequence. Statistical analyses of behavioral and FSCV data consisted of t tests and repeated-measures ANOVAs with post-hoc tests corrected for multiple comparisons as indicated throughout the corresponding figure legends. Stimulation-evoked dopamine traces also were analyzed with Difference Traces that digitally subtracted the Passive Playback response from the Self-Stimulation response for each pair of matched stimulations. Dopamine trace time courses following event onset were analyzed with permutation tests (10,000 random shuffles) with a cluster-based correction for multiple comparisons over time (Nichols & Holmes, 2002; Maris & Oostenveld, 2007). For electrophysiological data analysis, neuronal firing was aligned to stimulation onset, averaged within each session phase, and smoothed with a Gaussian filter (window size = 50 ms, standard deviation = 10) to construct peri-event time histograms for Self-Stimulation and Passive Playback responses. Statistical analyses were performed in Prism (GraphPad) and Matlab (MathWorks).

## ACKNOWLEDGMENTS

We thank Jared Smith, Jason Klug, Sho Aoki, Roy Kim, Kanchi Mehta, Anthony Balolong-Reyes, and Scott Ng-Evans for helpful discussions and technical assistance. This work was supported by NIH grants K99MH119312 (N.G.H.) and R01NS083815 (X.J.), the Salk Institute Pioneer Postdoctoral Endowment Fund and the Jonas Salk Fellowship (N.G.H.), and the McKnight Neurobiology of Brain Disorders Award (X.J.).

## ADDITIONAL INFORMATION

### FUNDING

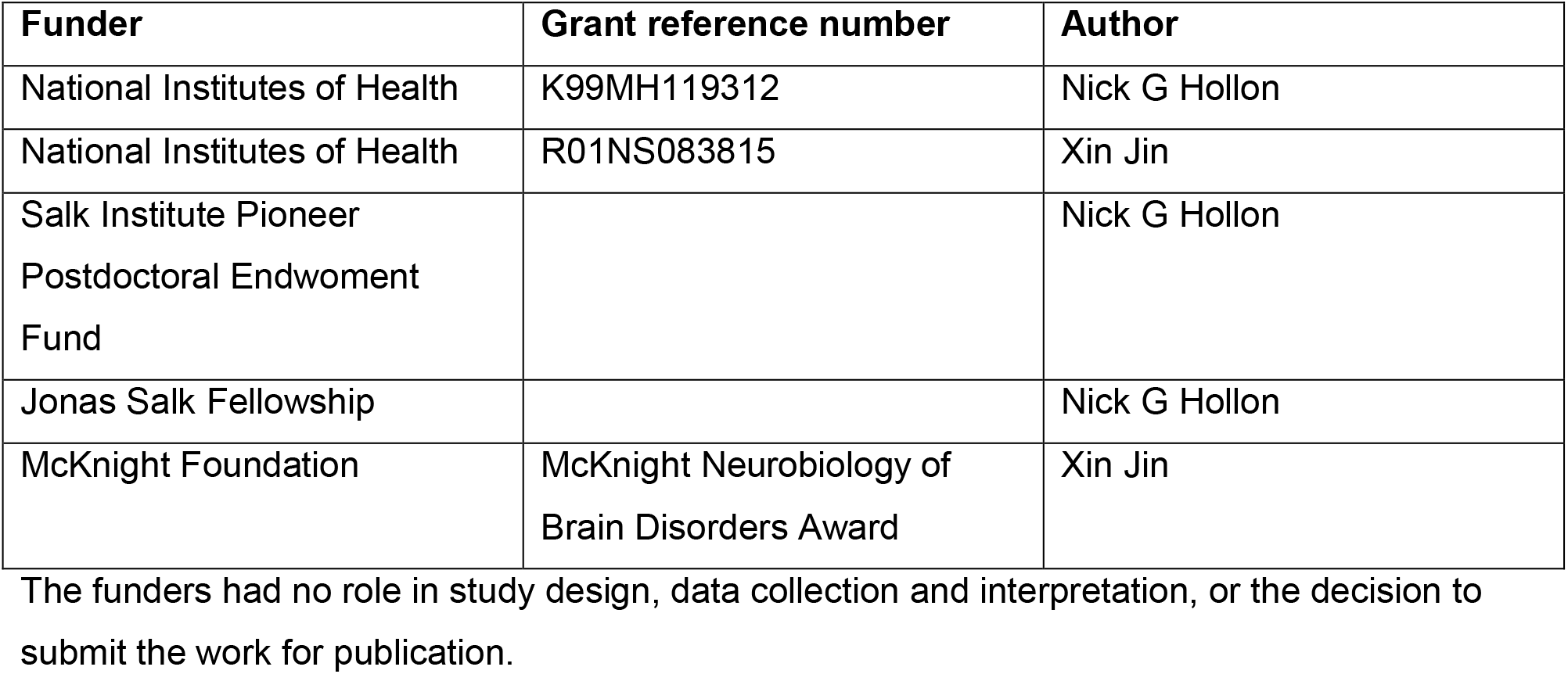

### AUTHOR CONTRIBUTIONS

Nick G Hollon, Conceptualization, Methodology, Software, Validation, Formal analysis, Investigation, Data curation, Writing—original draft, Writing—review and editing, Visualization, Supervision, Project administration, Funding acquisition; Elora W Williams, Validation, Formal analysis, Investigation, Data curation, Writing—review and editing; Christopher D Howard, Conceptualization, Methodology, Validation, Writing—review and editing, Supervision; Hao Li, Methodology, Software, Validation, Formal analysis, Investigation, Data curation, Writing—review and editing, Visualization; Tavish I Traut, Validation, Investigation, Writing— review and editing; Xin Jin, Conceptualization, Methodology, Validation, Resources, Writing— original draft, Writing—review and editing, Visualization, Supervision, Project administration, Funding acquisition

## SUPPLEMENTAL FIGURES

**Figure 1—figure supplement 1.**
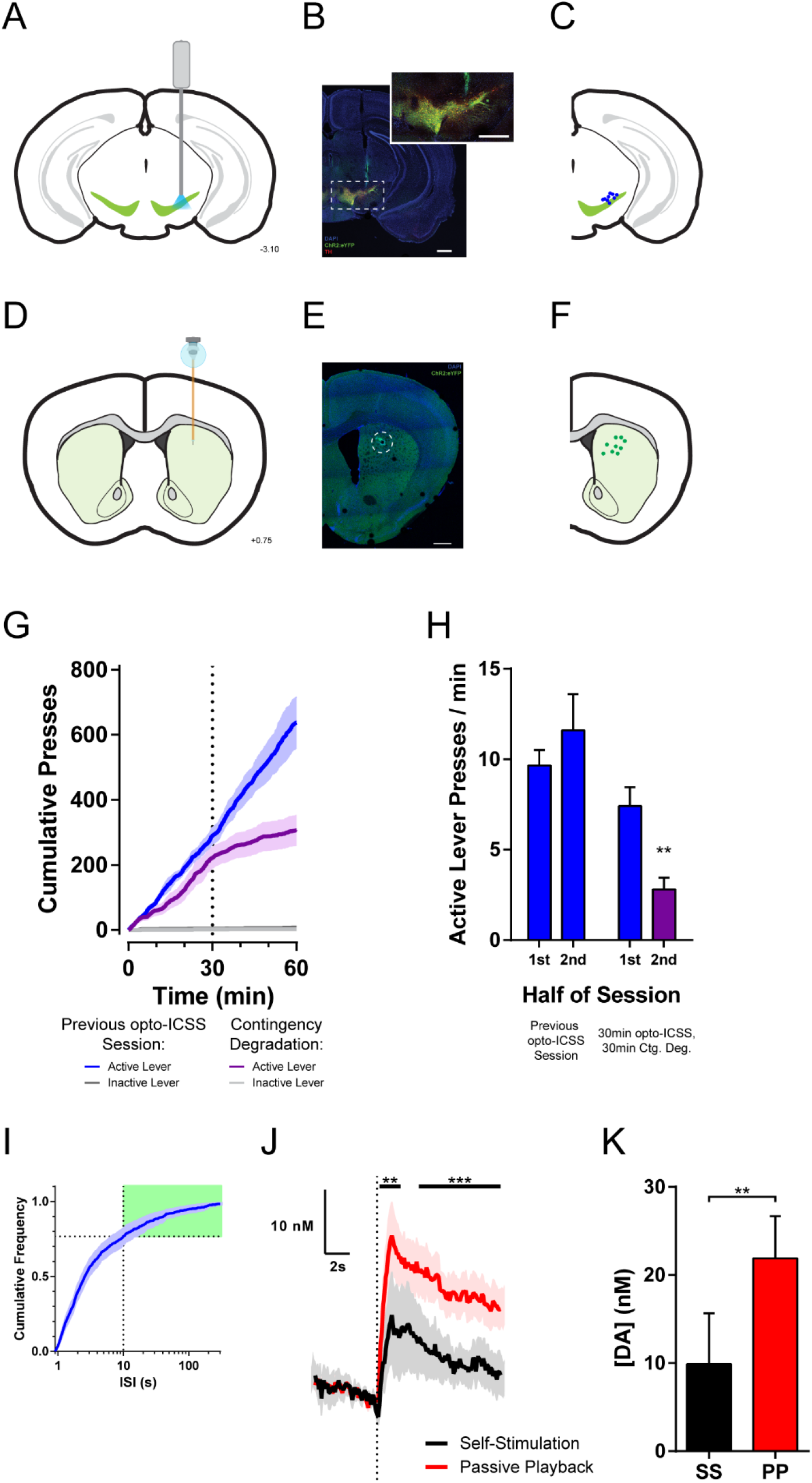
Opto-ICSS CRF Cohort Histology, Contingency Degradation, and Bout Initiations. (**A**) Coronal schematic of fiber optic placement targeting the SNc(−3.10 mm posterior from bregma). (**B**) Representative image of fiber optic placement over the SNc in animal selectively expressing ChR2-eYFP in TH-positive dopamine neurons (scale bars = 500 μm). (**C**) Fiber optic placement for mice in the CRF cohort. (**D**) Coronal schematic of FSCV carbon-fiber microelectrode placement in the dorsal striatum (+0.75 mm anterior to bregma). (**E**) Representative image of FSCV carbon-fiber microelectrode placement in the dorsal striatum (scale bar = 500 μm). The lesion was made by passing a current through the electrode just before perfusion after the conclusion of all experiments (see Methods). (**F**) FSCV electrode placement for mice in the CRF cohort. (**G**) Cumulative presses over time within the contingency degradation test session (30 min opto-ICSS followed by the 30 min contingency degradation test phase), overlaid with performance throughout the previous day’s standard opto-ICSS session for comparison (*n* = 6 mice). (**H**) Summary of mean Active lever press rate during each phase of the contingency degradation test session (as in Fig. 1A), compared to the preceding day’s standard opto-ICSS session (two-way repeated-measures ANOVA: main effect of Day, *F*_1,5_ = 8.157, *P* = 0.0356; Day by Half of Session interaction, *F*_1,5_ = 25.30, *P* = 0.0040; Sidak’s multiple comparisons tests: Contingency Degradation 1^st^ vs. 2^nd^ Half, *P* = 0.0082; 2^nd^ Half of Previous Day vs. Contingency Degradation test phase, *P* = 0.0004). (**I**) Cumulative frequency distribution of inter-stimulation intervals (ISIs) from the opto-ICSS FSCV recording session (*n* = 9 mice). Green shading indicates ISIs > 10 s, used to define bout initiation for the subset of stimulations analyzed in (J-K). (**J**) Mean dopamine concentration change to bout-initiating Self-Stimulation (ISI > 10s since previous stimulation) and corresponding Passive Playback stimulations. Black bars indicate time points where the Self-Stimulation response significantly differs from Passive Playback (permutation test, *P*s = 0.007 and 0.0001 for first and second time clusters, respectively). (**K**) Mean change in dopamine concentration for the bout-initiating subset of stimulations in (J). (*t*_8_ = 3.600, *P* = 0.0070). Ctg. Deg., Contingency Degradation; ISI, Inter-stimulation interval; SS, Self-Stimulation; PP, Passive Playback. Error bars are SEM here and for below figures.

**Figure 2—figure supplement 1.**
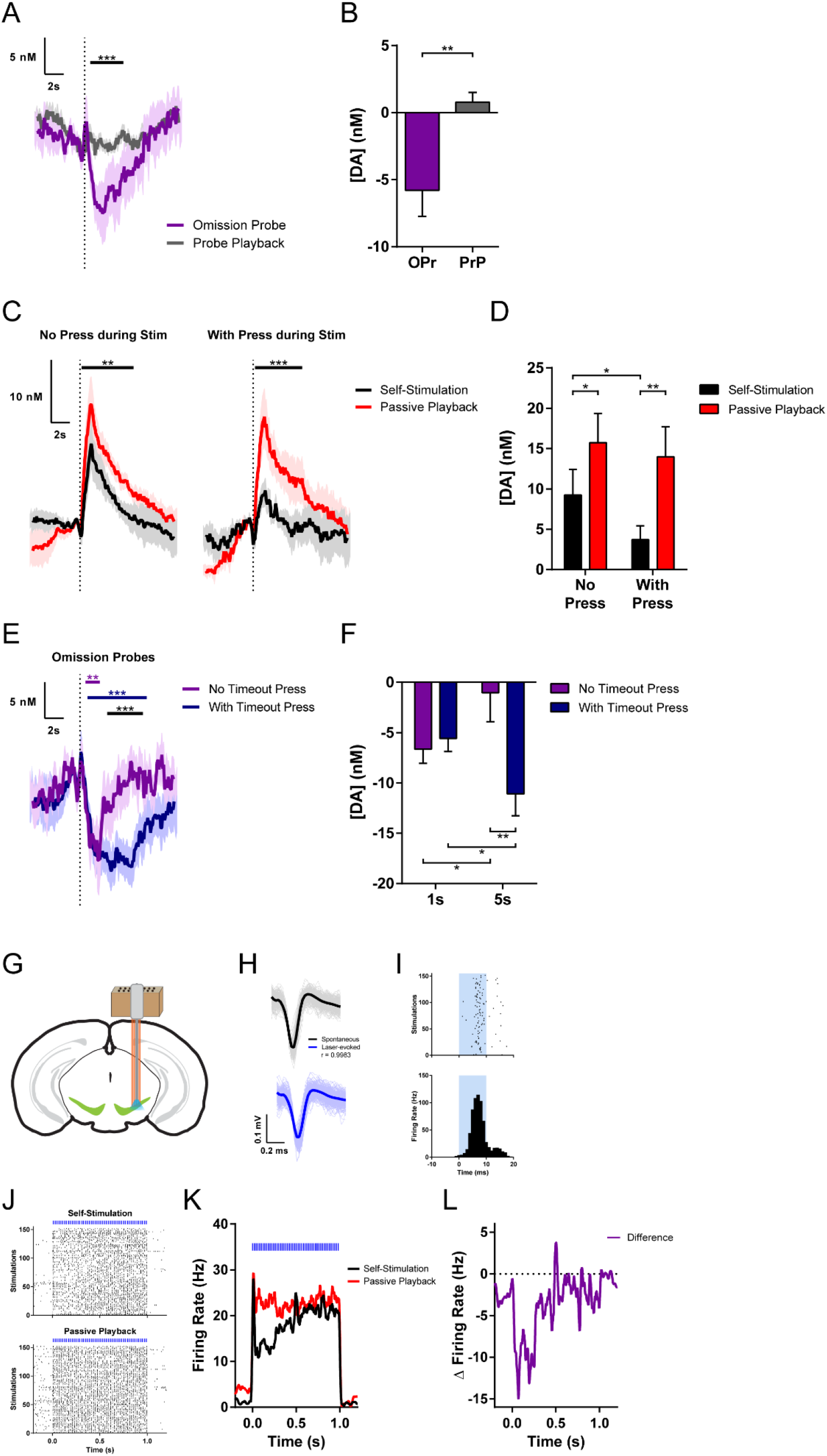
Inhibition of Dopamine to Isolated Omissions, Augmented Suppression by Additional Presses, and *In Vivo* **Electrophysiology**. (**A**) Mean dopamine concentration change following temporally isolated Omission Probes (latency > 5 s since previous stimulation) and corresponding time points from the Passive Playback phase (*n* = 8 mice; permutation test, *P* = 0.0005). (**B**) Mean change in dopamine concentration following temporally isolated Omission Probes and equivalent time points from Playback phase (paired t test, *t*_7_ = 3.511, *P* = 0.0098). (**C**) Mean dopamine concentration changes evoked by Self-Stimulations without (left) or with (right) an additional lever press during the ongoing stimulation, and the corresponding Passive Playback stimulations (*n* = 9 mice; permutation tests: Self-Stimulation with no press during stim vs. its Playback, *P* = 0.0011; Self-Stimulation with press during stim vs. its Playback, *P* = 0.0003; Self-Stimulation with vs. no press during stim, *P* = 0.0028). (**D**) Mean change in dopamine concentration for Self-Stimulations with or without additional presses during the stimulation, and their Playback (two-way repeated-measures ANOVA: main effect of Press, *F*_1,8_ = 8.144, *P* = 0.0214; main effect of Session Phase, *F*_1,8_ = 16.62, *P* = 0.0035; Sidak’s multiple comparisons tests: Self-Stimulation with vs. no additional press, *P* = 0.0350; Self-Stimulation without additional press vs. Playback, *P* = 0.0163; Self-Stimulation with additional press vs. Playback, *P* = 0.0011). (**E**) Mean dopamine concentration changes during Omission Probes with or without additional presses during the probe timeout period (*n* = 8 mice; permutation tests: Omission Probe with no timeout press vs. 0, magenta bar, *P* = 0.002; Omission Probe with timeout press vs. 0, blue bar, *P* = 0.0001; Omission Probe with vs. without press, black bar, *P* = 0.0003). (**F**) Mean change in dopamine concentration at early (1 s) vs. late (5 s) time points during Omission Probes with or without additional presses during the timeout period (two-way repeated-measures ANOVA: main effect of Press, *F*_1,7_ = 8.181, *P* = 0.0243; Press by Time interaction, *F*_1,7_ = 16.70, *P* = 0.0047; Sidak’s multiple comparisons tests: with vs. no press, late, *P* = 0.0025; no press, early vs. late, *P* = 0.0444; with press, early vs. late, *P* = 0.0479). (**G**) Schematic of experimental preparation for *in vivo* extracellular electrophysiology recordings with optogenetic identification of SNc dopamine neurons. (**H**) Waveforms of optogenetically identified dopamine neuron for spontaneous (top) and laser-evoked (bottom) spikes (Pearson correlation, r = 0.9983, *P* < 0.0001). (**I**) Raster plot (top) and peri-event time histogram (bottom) of dopamine neuron response to 10-ms optogenetic stimulation pulse. Each row in the raster represents one stimulation, and black ticks are spikes. (**J**) Raster plot of the same dopamine neuron responses aligned to Self-Stimulation (top) and Passive Playback stimulations (bottom). (**K**) Firing rate of the optogenetically identified dopamine neuron in (J) in response to Self-Stimulation versus Passive Playback stimulations. (**L**) Difference trace for the dopamine neuron in (J) depicting Self-Stimulation minus Playback difference in stimulation-evoked firing rate between session phases. OPr, Omission Probe; PrP, Probe Playback.

**Figure 3—figure supplement 1.**
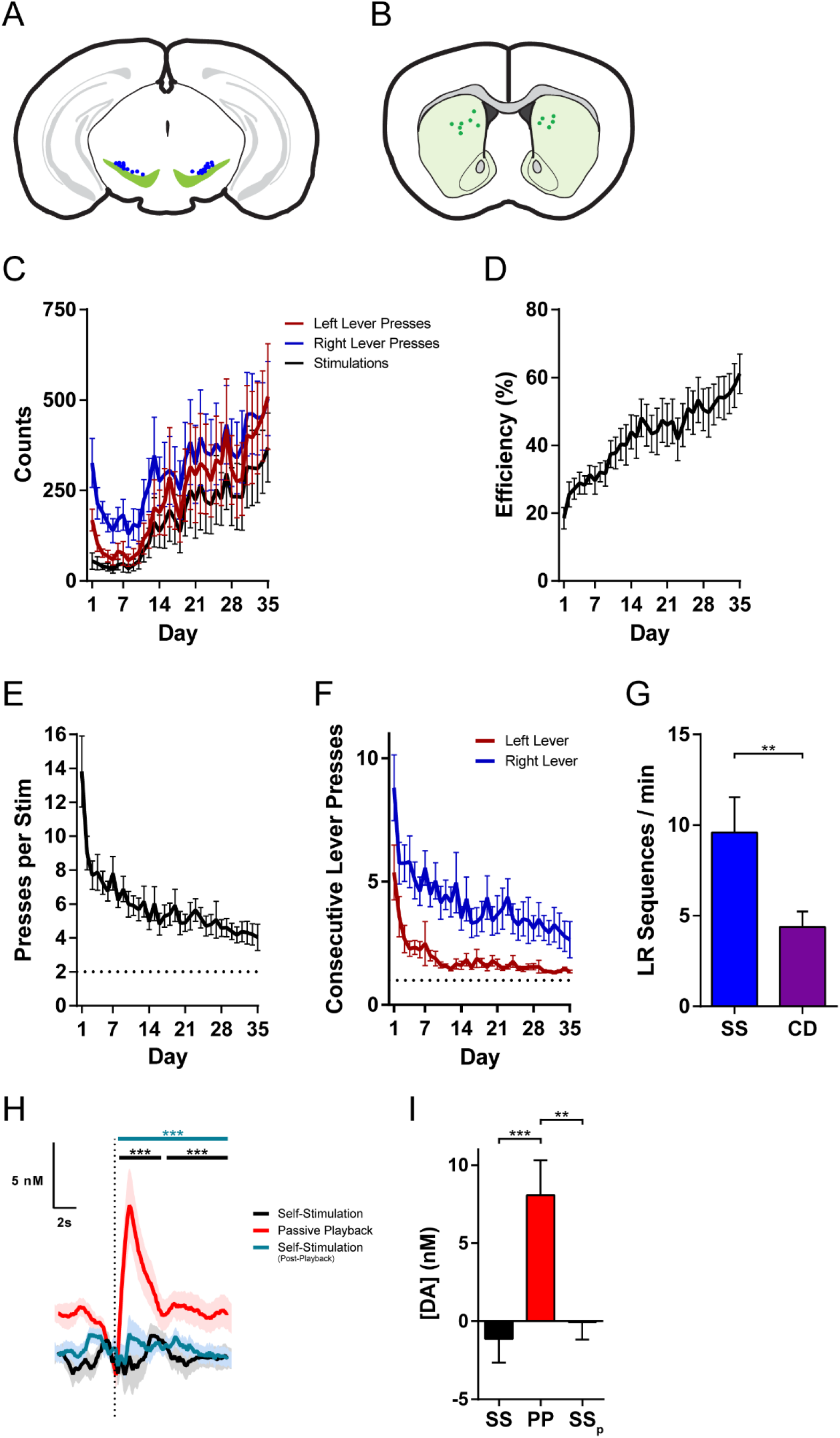
Left-Right Sequence Cohort Histology, Additional Behavior, and Session Phase Order Control. (**A**) Fiber optic placement for mice in the LR sequence cohort. (**B**) FSCV electrode placement for mice in the LR sequence cohort. (**C**) Presses on each lever and stimulations earned across days of training (*n* = 13 mice; statistics for stimulations presented in Fig. 3B; two-way repeated-measures ANOVA for Lever by Day: main effect of Day, *F*_34,408_ = 3.189, *P* < 0.0001; main effect of Lever, *F*_1,12_ = 16.67, *P* = 0.0015; Lever by Day interaction, *F*_34,408_ = 1.535, *P* = 0.0307). (**D**) Efficiency across days of training, calculated as the number of stimulations per pair of lever presses (one-way repeated-measures ANOVA, *F*_34,408_ = 6.936, *P* < 0.0001). (**E**) Total presses per stimulation (either lever) across days of training (one-way repeated-measures ANOVA, *F*_34,408_ = 8.122, *P* < 0.0001). (**F**) Consecutive presses on each lever across days of training (two-way repeated-measures ANOVA: main effect of Day, *F*_34,408_ = 8.430, *P* < 0.0001; main effect of Lever, *F*_1,12_ = 23.07, *P* = 0.0004). (**G**) Contingency degradation test: 30 min of LR sequence opto-ICSS followed by 30 min contingency degradation test phase (n = 10 mice; paired t test, *t*_9_ = 3.458, *P* = 0.0072). (**H**) Mean dopamine concentration change evoked by LR Self-Stimulation before or after the Passive Playback phase. The pre-playback Self-Stimulation (black) and Passive Playback responses are the same data as Fig. 3H, and the post-playback Self-Stimulation (teal) is an additional 30 min phase of LR sequence opto-ICSS following the Playback phase to control for potential order effects (*n* = 12 mice; permutation tests: *P*s = 0.0001 for all time clusters, black bars for pre-playback Self-Stimulation vs. Playback, teal bar for post-playback Self-Stimulation vs. Playback). (**I**) Mean change in dopamine concentration for Self-Stimulation before or after Passive Playback (one-way repeated-measures ANOVA, *F*_2,22_ = 26.39, *P* < 0.0001; Tukey’s multiple comparisons tests: Self-Stimulation (pre) vs. Playback, *P* < 0.0001; Self-Stimulation (post) vs. Playback, *P* < 0.0001). SS, Self-Stimulation; CD, Contingency Degradation; PP, Passive Playback; SSP, Self-Stimulation (post-Playback).

